# Characterization of cefotaxime resistant *Escherichia coli* isolated from broiler farms in Ecuador

**DOI:** 10.1101/462994

**Authors:** Christian Vinueza-Burgos, David Ortega-Paredes, Cristian Narváez, Lieven De Zutter, Jeannete Zurita

## Abstract

Antimicrobial resistance (AR) is a worldwide concern. Up to a 160% increase in antibiotic usage in food animals is expected in Latin American countries. The poultry industry is an increasingly important segment of food production and contributor to AR. The objective of this study was to evaluate the prevalence, AR patterns and the characterization of relevant resistance genes in Extended Spectrum β-lactamases (ESBL) and AmpC *E. coli* from large poultry farms in Ecuador. Sampling was performed from June 2013 to July 2014 in 6 slaughterhouses that slaughter broilers from 115 farms totaling 384 flocks. Each sample of collected caeca was streaked onto TBX agar supplemented with cefotaxime (3 mg/l). In total, 176 isolates were analyzed for antimicrobial resistance patterns by the disk diffusion method and for *bla*_CTX-M_, *bla*_TEM_, *bla*_CMY_, *bla*_SHV_, bla_*KPC*_, and *mcr-1* by PCR and sequencing. ESBL and AmpC *E. coli* were found in 362 flocks (94.3%) from 112 farms (97.4%). We found that 98.3% of the isolates were multi-resistant to antibiotics. Low resistance was observed for ertapenem and nitrofurantoin. The most prevalent ESBL genes were the *bla_CTX-M_* (90.9%) *bla_CTX-M-65_*, *bla_CTX-M-55_* and *bla_CTX-M-3_* alleles. Most of the AmpC strains presented the *bla_CMY-2_* gene. Three isolates showed the *mcr-1* gene. Poultry production systems represent a hotspot for antimicrobial resistance in Ecuador, possibly mediated by the extensive use of antibiotics. Monitoring this sector in national and regional plans of antimicrobial resistance surveillance should therefore be considered.

## Introduction

Antimicrobial resistance (AR) is a worldwide concern. It is expected that deaths linked to antimicrobial resistance could rise from 700,000 to 10 million deaths per year by 2050[1]. On the other hand, 23.000 (United States) to 25.000 (Europe) deaths could be attributed to resistant pathogens in developed countries [2].

In terms of economic loses, AR could cause a global loss of production as high as 100 trillion dollars which represents a huge impact on the economy of all countries, especially developing countries [1]. This problem will not only result in increased illnesses, disabilities and deaths but also puts at risk the achievement of Sustainable Development Goals for the next 30 years [3,4].

It has been observed that AR in relevant pathogens has increased in recent years [5,6]. The causes of this phenomenon are complex and are mainly linked to selective pressures triggered by antibiotic usage [7]. Inappropriate prescription of antimicrobials, unauthorized sale of antibiotics and the extensive use of these drugs in animal production are some factors that contribute to this problem [8,9]. For example, in 2010, more than 63.000 tons of antibiotics were used in livestock [10]. Moreover, it has been projected that by 2030 the use of antibiotics in livestock will double in countries such as Brazil, Russia, India and South Africa. Furthermore, up to a 160% increase in antibiotic usage in food animals is expected in Latin American countries [11].

Worldwide, the poultry industry is an increasingly important segment of food production. In fact, it is expected that by 2025, poultry will be the most important sector of meat production [11,12]. Widespread use of antibiotics in the poultry industry impacts the AR problem. This issue is especially relevant in developing countries where antimicrobials are not only used therapeutically but also prophylactically and as growth promoters [13,14].

Among the antibiotics used in livestock production, some are listed in the WHO list of critically important antimicrobials for human medicine. This group includes third generation cephalosporines, carbapenems and colistin, all of which are categorized as highest priority [15]. Additionally, ESBL- and AmpC-producing *E. coli*, and carbapenem-resistant *E*. *coli* are listed as high priority organisms for which new antibiotics are urgently needed [16].

*E. coli* harboring resistance determinants originating in the poultry industry are therefore of great epidemiological interest because they can serve as reservoirs of resistance genes that can be transferred to human pathogens [17]. A relationship between resistant strains of *E. coli* from poultry and those found in humans has been suggested in several studies [18-20]. However, information about resistant *E. coli* in industrial poultry has been poorly studied in Latin America. The objective of this study was to evaluate the prevalence and AR patterns of and to characterize relevant resistance genes in ESBL and AmpC *E. coli* from large poultry farms in Ecuador.

## Material and methods

### Study Design and Sampling

Pichincha, the province where Quito, the capital city of Ecuador is located, was selected for the collection of samples since 36% of the total Ecuadorian broiler production is located in this and surrounding provinces [21]. Eight large slaughterhouses are located in Pichincha [21] and all were asked to participate in the study. Sampling was performed in the 6 participating slaughterhouses which slaughter broilers from 115 farms. From June 2013 to July 2014, a total of 384 flocks (birds coming from one house and slaughtered on the same day) were sampled. All sampled flocks from the same farm originated from different houses or birds reared during different periods in the same house.

In Ecuador, commercial broiler farm management includes the total depopulation of houses and removal of the litter after every flock, cleaning and disinfection of the house followed by a dormant period of 8 to 15 days. All sampled flocks were commercially reared and slaughtered at the age of 6 to 7 weeks. From each batch, caeca from 25 randomly selected chickens were collected”, and transported in an ice box within 1 hour to the laboratory for bacteriological analysis.

### Isolation and Identification of ESBL/AmpC *E. coli*

Caeca from each flock were immersed in 98% ethanol, and after evaporation of the ethanol, approximately 1 g of fecal content was collected and pooled in a sterile plastic bag. The pooled sample was homogenized by hand for 1 minute.

Each sample was streaked onto TBX agar (BioRad) supplemented with cefotaxime (3 mg/l). Two sample colonies were confirmed using Triple Sugar Iron agar (Difco, BD). From this media, one loopful was used to extract DNA by the boiling method. Another loopful was used to subculture the isolate in trypticase soy broth (Difco, BD) and stored with glycerol (60%) at −80°C. All cefotaxime resistant *E. coli* isolates were further examined for the presence of ESBL using ceftazidime, ceftazidime/clavulanate, cefotaxime, cefotaxime/clavulanate disks [22] and for the AmpC phenotype using boronic acid, ceftazidime and cefepime disks [23].

### Antimicrobial Resistance and PCR screening

One isolate with the ESBL and AmpC phenotypes from each farm was selected for analysis by the Kirby Bauer method. Antimicrobial resistance profiles were evaluated using clinical breakpoint values from the Clinical and Laboratory Standards Institute [22]. The following antibiotics were evaluated: trimethoprim-sulfamethoxazole, nalidixic acid, ciprofloxacin, gentamicin, kanamycin, streptomycin, tetracycline, chloramphenicol, fosfomycin, tetracycline, doxycycline, ceftazidime and ertapenem. *E. coli* ATCC 25922 was used as a quality control strain.

Selected isolates were studied by PCR to identify *bla*_CTX-M_, *bla*_TEM_, *bla*_CMY_, *bla*_SHV_ and bla*_KPC_*. PCR conditions and primers were those described by [24] for *bla*_CTX-M_, [25] for *bla*_TEM_, [26] for *bla*_CMY_, [27] for *bla*_SHV_ and [28] for bla_*KPC*_. Sub-families of *bla*_CTX-M_ genes were identified with PCR protocols described by [29] for *bla*_CTX-M-1_, [30] for *bla*_CTX-M-2_, [31] for *bla*_CTX-M-8_, [32] for *bla*_CTX-M-9_ and [33] for *bla*_CTX-M-14-like_. Isolates with phenotypic resistance to colistin were tested for the presence of *mcr*-1 [34] and *mcr*-2 plasmid genes [35]. Amplification products were confirmed by gel electrophoresis using a 1% agarose gel. All PCR products were purified and sequenced at Macrogen Inc. (Seoul, South Korea). Obtained sequences were aligned against reference sequences with the online tool ResFinder 2.1 [36]. MIC values for ampicillin, tazobactam, cefoxitin, ceftazidime, ceftriaxone, cefepime, doripenem, ertapenem, imipenem, meropenem, amikacin, gentamicin, ciprofloxacin, tigecycline and colistin were obtained on *mcr*-1 positive isolates using the Vitek 2 system with the AST-N272 card. The results were evaluated using the breakpoints recommended by CLSI [22].

### Genetic characterization

Fingerprint characterization was performed in selected isolates by repetitive element palindromic PCR (REP-PCR) analysis [37]. Bands were analyzed using BioNumerics software V.7.6 (Applied Maths, Sint-Martems-Latem, Belgium). Fragments between 200 bp and 1500 bp in size were included in the analysis. The unweighted pair group method using the arithmetic averages algorithm (UPGMA) with a 1.5% tolerance was used to construct a dendrogram.

### Statistical Analysis

To determine the prevalence of *E. coli* ESBL at the farm level, a farm was considered positive when at least one of the sampled batches was positive. Farms were assumed to be independent.

Differences in antibiotic resistances between ESBL *E. coli* and AmpC *E. coli* were calculated by the chi-square test. Proportions were considered significantly different when the *P* value was below 0.05.

## Results

### Prevalence of ESBL and AmpC *E. coli* isolates at Poultry Farms

ESBL and AmpC *E. coli* were found in 362 flocks (94.3%; CI95%: 93,39% – 95,26%). In total, 112 farms (97.4%; CI95%: 96.37% – 98.94%) delivered a positive result at least once.

From all positive flocks, 62 (17.1%) delivered a combination of ESBL and AmpC isolates while 223 (61.6%) and 51 (14.1) flocks had exclusively the ESBL or AmpC phenotypes, respectively. Only one colony could be isolated from 26 flocks, 21 (5.8%) and 3 (0.01%) showing the ESBL and AMPC phenotypes respectively, and 2 flocks produced no colonies. For antimicrobial resistance tests, 110 *E. coli* ESBL and 66 *E. coli* AmpC isolates were selected for further analysis.

### Antimicrobial resistance patterns

Antimicrobial susceptibility testing grouped ESBL and AmpC *E. coli* isolates in 26 patterns that showed resistances to between 2 and 7 antibiotic families tested. Resistance patterns to at least 3 antibiotic families (multi-resistant isolates) were present in 98.3% of all tested isolates. Moreover, 92.1% of isolates presented resistance to between 4 and 7 antibiotic families. Pattern number 3 was the most common one for ESBL isolates with resistance to all tested groups of antibiotics with the exception of nitrofurantoin and pattern number 1 for AmpC isolates with resistance to all tested groups of antibiotics (Table 1). Significantly, 9.7% of ESBL or AmpC *E. coli* isolates presented resistance to all tested groups of antibiotics.

**Table 1.** Antibiotic resistance patterns of ESBL/AmpC *E. coli* isolated from poultry farms.

| Pattern | Resistance Pattern number | No. of antibiotic groups | No. of ESBL isolates | No. of AmpC isolates | Total isolates (%) |
| --- | --- | --- | --- | --- | --- |
| BQTAPNS | 1 | 7 | 8 | 9 | 17 (9.7) |
| BQTANS | 2 | 6 | 2 | 2 | 4 (2.3) |
| BQTAPS | 3 | 6 | 42 | 19 | 61 (34.7) |
| BQTAPN | 4 | 6 | 1 | 2 | 3 (1.7) |
| BQTPNS | 5 | 6 | 1 | 1 | 2 (1.1) |
| BQTAS | 6 | 5 | 9 | 6 | 15 (8.5) |
| BQTAN | 7 | 5 | 1 | 1 | 2 (1.1) |
| BQTNS | 8 | 5 | 1 |  | 1 (0.6) |
| QTAPS | 9 | 5 |  | 4 | 4 (2.3) |
| BQTAP | 10 | 5 | 6 | 3 | 9 (5.1) |
| BTAPS | 11 | 5 | 1 | 2 | 3 (1.7) |
| BQAPS | 12 | 5 | 4 | 3 | 7 (4) |
| BAPNS | 13 | 5 | 1 |  | 1 (0.6) |
| BQTPS | 14 | 5 | 2 | 1 | 3 (1.7) |
| BQTA | 15 | 4 | 12 | 1 | 13 (7.4) |
| BQTS | 16 | 4 |  | 2 | 2 (1.1) |
| BQAN | 17 | 4 |  | 1 | 1 (0.6) |
| BAPS | 18 | 4 | 2 | 1 | 3 (1.7) |
| BQPS | 19 | 4 | 1 | 1 | 2 (1.1) |
| BQAP | 20 | 4 | 3 | 2 | 5 (2.8) |
| BQTP | 21 | 4 | 1 | 1 | 2 (1.1) |
| BTPS | 22 | 4 | 2 |  | 2 (1.1) |
| BQT | 23 | 3 | 5 | 1 | 6 (3.4) |
| BQA | 24 | 3 | 1 | 1 | 2 (1.1) |
| BTA | 25 | 3 | 1 | 2 | 3 (1.7) |
| BQ | 26 | 2 | 3 |  | 3 (1.7) |
| **TOTAL** |  |  |  |  | **110** |

Antimicrobial resistance rates for ESBL and AmpC *E. coli* isolates are shown in Table 2. Low resistance rates were observed for ertapenem followed by nitrofurantoin. For the remaining antibiotics, resistance rates ranged from 29,1% to 93.9%. Antibiotics for which significant differences were observed between ESBL and AmpC isolates were ceftazidime, kanamycin and gentamicin. AmpC isolates presented phenotypes with higher resistance to ceftazidime and kanamycin while ESBL isolates were more resistant to gentamicin (Table 2).

**Table 2.**
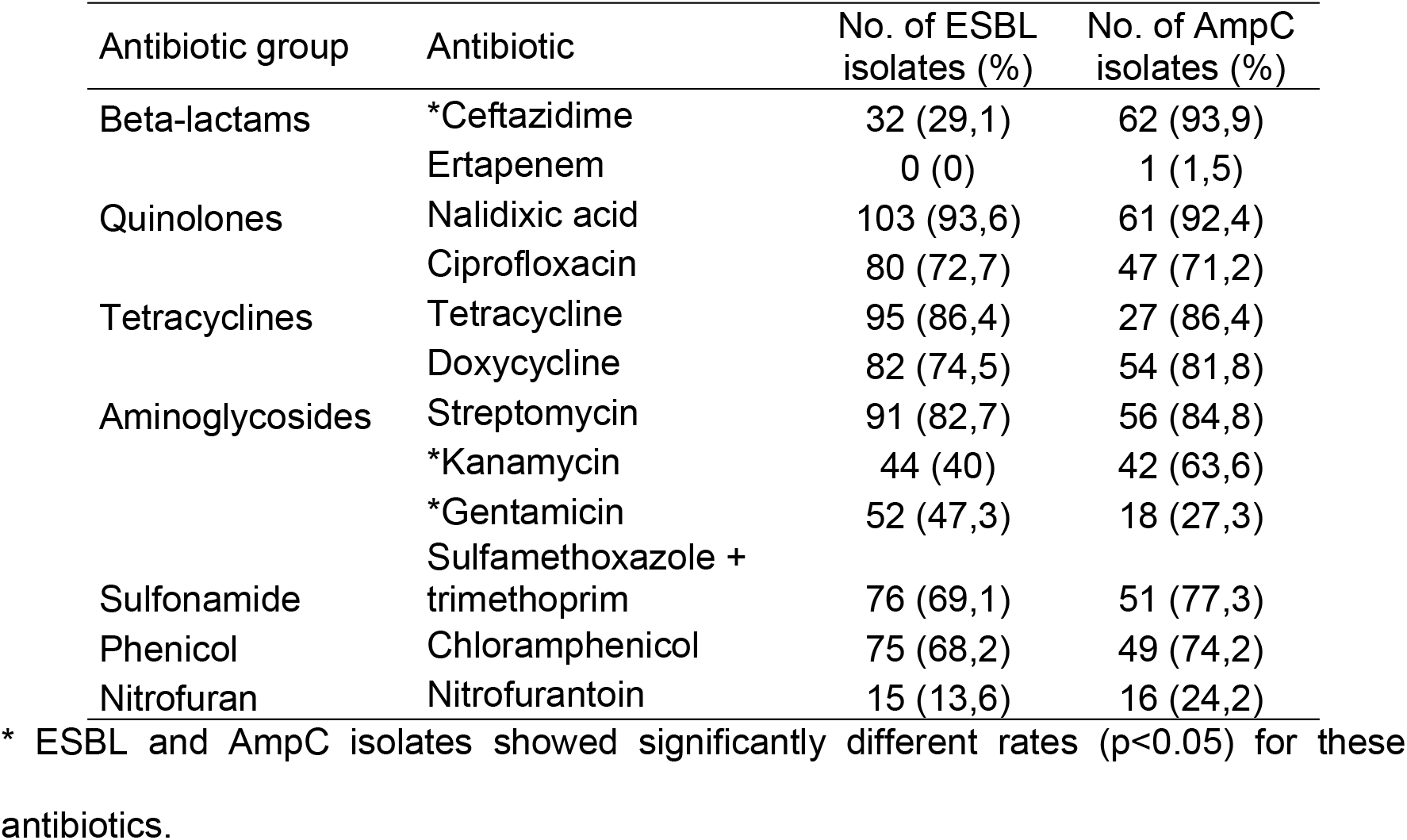
Number of ESBL/AmpC *E. coli* isolates resistant to each tested antibiotic.

Three isolates from the ESBL group and 3 isolates from the AmpC group were positive for the *mcr-*1 gene. MIC values of these isolates are shown in Table 3. All these isolates presented resistance to ceftriaxone, and colistin. Resistance to doripenem, imipenem, amikacin and tigecycline was not identified in any isolate. Two and 3 isolates were resistant to ceftazidime and cefoxitin respectively. Only isolates 1CT22A, 1CT86A and 1CT160A presented phenotypical resistance to piperacillin-tazobactam, cefepime and, meropenem and ertapenem respectively.

**Table 3.**
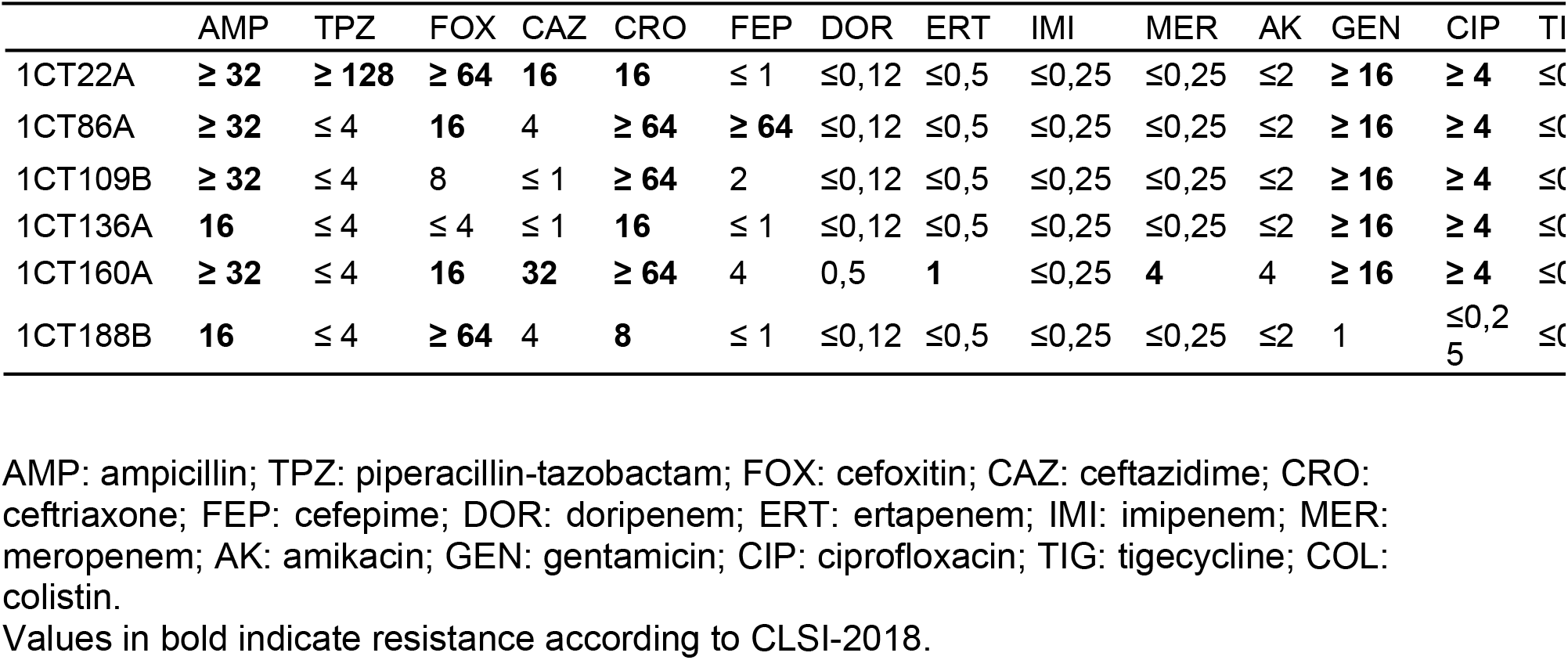
MIC values of *mcr*-1 positive isolates.

|  | AMP | TPZ | FOX | CAZ | CRO | FEP | DOR | ERT | IMI | MER | AK | GEN | CIP | TIG | COL |
| --- | --- | --- | --- | --- | --- | --- | --- | --- | --- | --- | --- | --- | --- | --- | --- |
| 1CT22A | ≥ 32 | ≥ 128 | ≥ 64 | 16 | 16 | ≤ 1 | ≤ 0,12 | ≤ 0,5 | ≤ 0,25 | ≤ 0,25 | ≤ 2 | ≥ 16 | ≥ 4 | ≤ 0,5 | 8 |
| 1CT86A | ≥ 32 | ≤ 4 | 16 | 4 | ≥ 64 | ≥ 64 | ≤ 0,12 | ≤ 0,5 | ≤ 0,25 | ≤ 0,25 | ≤ 2 | ≥ 16 | ≥ 4 | ≤ 0,5 | 8 |
| 1CT109B | ≥ 32 | ≤ 4 | 8 | ≤ 1 | ≥ 64 | 2 | ≤ 0,12 | ≤ 0,5 | ≤ 0,25 | ≤ 0,25 | ≤ 2 | ≥ 16 | ≥ 4 | ≤ 0,5 | 2 |
| 1CT136A | 16 | ≤ 4 | ≤ 4 | ≤ 1 | 16 | ≤ 1 | ≤ 0,12 | ≤ 0,5 | ≤ 0,25 | ≤ 0,25 | ≤ 2 | ≥ 16 | ≥ 4 | ≤ 0,5 | 8 |
| 1CT160A | ≥ 32 | ≤ 4 | 16 | 32 | ≥ 64 | 4 | 0,5 | 1 | ≤ 0,25 | 4 | 4 | ≥ 16 | ≥ 4 | ≤ 0,5 | ≥ 16 |
| 1CT188B | 16 | ≤ 4 | ≥ 64 | 4 | 8 | ≤ 1 | ≤ 0,12 | ≤ 0,5 | ≤ 0,25 | ≤ 0,25 | ≤ 2 | 1 | ≤ 0,25 | ≤ 0,5 | 8 |

AMP: ampicillin; TPZ: piperacillin-tazobactam; FOX: cefoxitin; CAZ: ceftazidime; CRO: ceftriaxone; FEP: cefepime; DOR: doripenem; ERT: ertapenem; IMI: imipenem; MER: meropenem; AK: amikacin; GEN: gentamicin; CIP: ciprofloxacin; TIG: tigecycline; COL: colistin.
Values in bold indicate resistance according to CLSI-2018.

### Genetic characterization

Sequencing of ESBL genes showed that the most prevalent family of genes was *bla_CTX-M_* (90.9%). Within this family, *bla_CTX-M-65_*, *bla_CTX-M-55_* and *bla_CTX-M-3_* accounted for 70% of isolates. Forty-eight (43.6%) and 9 (8.2%) *bla_CTX-M_* positive isolates presented the *bla_TEM-1A_* or *bla_TEM-1B_*, and *bla_SHV-5_* alleles respectively. Three isolates had the *bla_TEM-176_* gene and 1 isolate presented the *bla_SHV-153_* gene. One and 4 isolates presented only *bla*_TEM-1A_ or *bla*_TEM-1B_ genes respectively which do not hydrolyze cefotaxime. One isolate was not positive for any of the studied ESBL genes (Table 4).

**Table 4.**
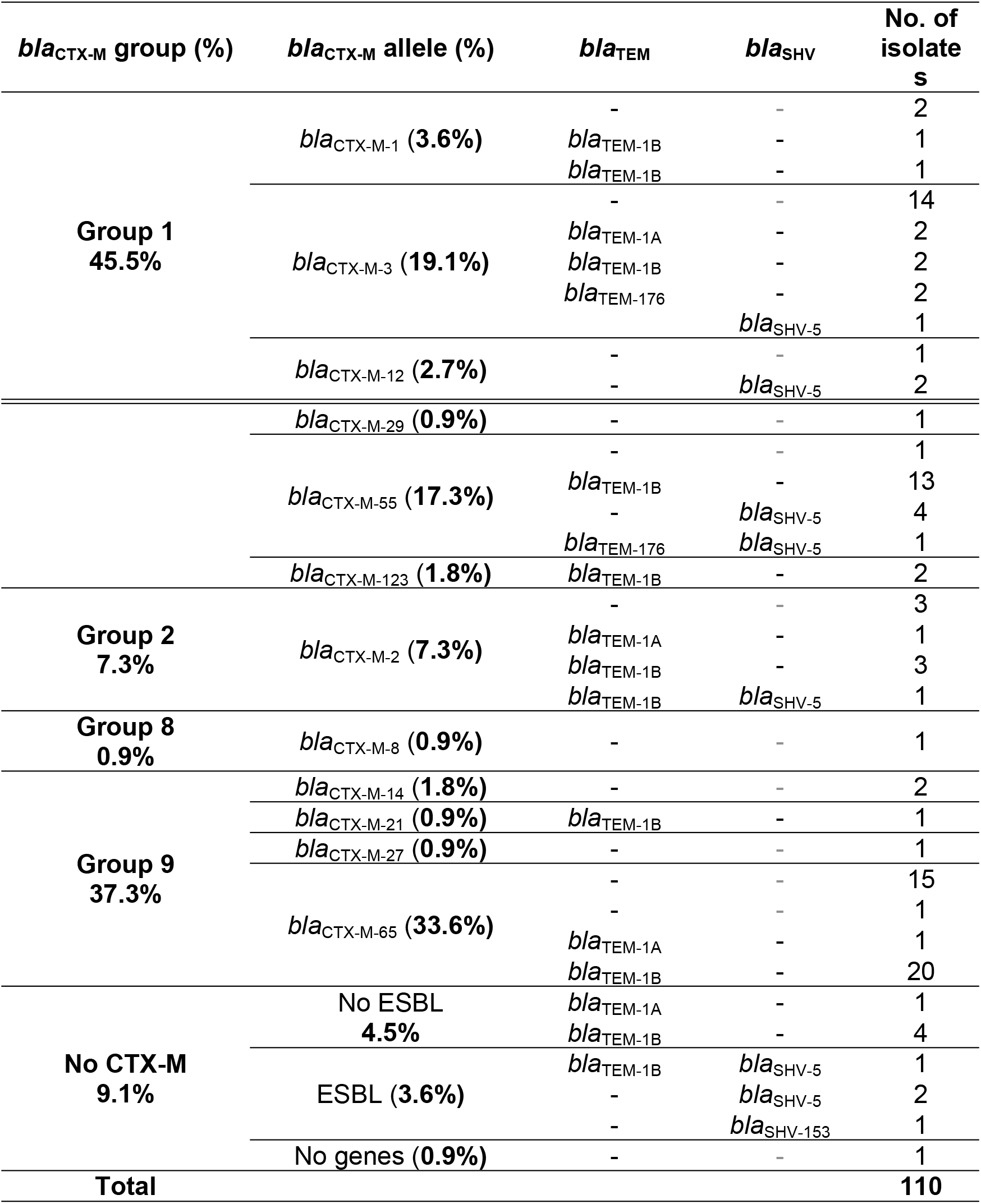
Combination of *bla_CTX-M_*, *bla_TEM_* and *bla_SHV_* alleles in ESBL *E. coli* isolates.

Of the 66 AmpC isolates, 7 were PCR negative for *bla_cmy_*, 58 were positive for *bla_cmy-2_*, and 1 isolate was positive for *bla_cmy-46_*. Each of the 3 Isolates from the ESBL group that were positive for the *mcr-1* gene showed the common presence of *bla_TEM-1B_*/*bla_CTX-M-1_*, *bla_SHV-5_*/*bla_CTX-M-55_* and *bla_CTX-M-65_* respectively. On the other hand, all *mcr-1-*positive isolates from the AmpC group were positive for *bla_cmy-2_*. None of the tested isolates were positive for *bla_KPC_* or *mcr*-2.

REP-PCR delivered 121 genotypes from which 22 grouped to more than one isolate (Table 5).

**Table 5.** Genotypes of ESBL and AmpC *E. coli* with more than one isolate.

| Genotypes | ESBL | AmpC | Total |
| --- | --- | --- | --- |
| I | 2 |  | 2 |
| II | 2 |  | 2 |
| III | 1 | 1 | 2 |
| IV | 2 |  | 2 |
| V |  | 2 | 2 |
| VI | 14 |  | 14 |
| VII | 3 | 2 | 5 |
| VIII |  | 2 | 2 |
| IX | 1 | 1 | 2 |
| X | 2 |  | 2 |
| XI | 10 | 1 | 11 |
| XII |  | 2 | 2 |
| XIII |  | 3 | 3 |
| XIV | 4 | 1 | 5 |
| XV | 2 |  | 2 |
| XVI | 4 |  | 4 |
| XVII | 2 |  | 2 |
| XVIII |  | 2 | 2 |
| XIX |  | 2 | 2 |
| XX | 3 |  | 3 |
| XXI | 3 |  | 3 |
| XXII | 2 |  | 2 |

Genotypes III, VII, IX, XI and XIV were common to ESBL and AmpC isolates. In total, genotypes VI and XI were the most common ones, with 14 and 11 isolates, respectively. Isolates from each genotype originated from different farms.

## Discussion

The aim of this research was to study antimicrobial resistance in *E. coli* from intensive poultry farming. The lack of this kind of data from Latin America makes this study one of the few available reports that demonstrate the extent of ESBL/AmpC *E. coli* in commercial poultry in the region [38-40]. Nonetheless, developed countries have also reported both a high prevalence of ESBL *E. coli* and the presence of multiresistant isolates from broiler flocks [41,42].

Similar to other studies, this research shows a higher prevalence of ESBL genes than AmpC genes. However, a study carried out in Colombia by Castellanos et al. [39] demonstrates a higher prevalence of AmpC genes in *E. coli* isolated from commercial poultry. Differences in the epidemiologic patterns of enteric bacteria isolated from Ecuadorian and Colombian poultry has been reported before and may be attributed to the ecological characteristics of the boundary between these countries [13].

High antimicrobial resistance rates and multi-resistance patterns could be related to the intensive use of antimicrobials in poultry production, which in some cases are not only used as therapeutics but also as prophylactics and growth promotors [11,43]. On the other hand, it has to be considered that the withdrawal of antibiotics from poultry production systems may not result in the diminution of ESBL/AmpC *E. coli* since ecological factors could be implicated in the dynamics of AR determinants [44,45].

Increasing antibiotic resistance and the lack of new antibacterial agents have revived interest in old compounds such as nitrofurantoin in clinical [46,47]. Despite the renewed importance of nitrofurantoin and the known role of food animals in resistance dissemination, only a few studies include this antibiotic in antimicrobial resistance screenings [48,49].

Resistance rates to nitrofurantoin in this study are higher than the ones reported in chicken meat samples (7,9%) in Colombia by Donado-Godoy et al. [49]. Higher resistance to nitrofurantoin in extra-intestinal clinical isolates of *E. coli* from chickens has been reported in China [48]. Due to the possibility that the resistance to nitrofurantoin could increase over time, it is important to include it in antimicrobial resistance surveillance plans in poultry production systems.

Carbapenems are not commonly used in the poultry industry, resulting in low selective pressure by this antimicrobial in poultry production. This observation explains the low prevalence of carbapenem-resistant *E. coli* reported in poultry [50]. Concordantly, we did not identify any isolate resistant to ertapenem, although carbapenem resistance mediated by *bla*_KPC_ has been reported in clinical Enterobacteriaceae in Ecuador [51].

The association of ESBL and AmpC phenotypes with increased prevalence of aminoglycoside resistance has been reported [52]. In our study we identified a significant association of kanamycin resistance with AmpC-producing isolates and gentamicin resistance to ESBL-producing *E. coli*. Further genetic characterization should be performed in order to explain whether these antimicrobial resistances are associated with specific genetic environments (the presence of ESBL/AmpC enzymes), In our study, all selected isolates were resistant to cefotaxime (used for screening), however, AmpC isolates were significantly more resistant to ceftazidime than ESBL isolates. This difference is explained by the enhanced hydrolyzation of ceftazidime by the *bla_CMY_* gene product [53,54].

Several studies throughout the world have reported plasmid-mediated colistin resistance in Enterobacteriaceae pointing to its global emergence [55]. In our study 6 out of 176 isolates (3.4%) were PCR positive for *mcr-1* and confirmed as phenotypically colistin resistant. In contrast, a study in Argentina reported that 49% (n=304) of *E. coli* isolates recovered from broilers were identified as colistin resistant by microdilution [56]. Another study in Brazil reported that 19.5% of chicken meat samples (n=41) were positive for *mcr*-1 [57]. Colistin resistant Enterobacteriaceae have been described in humans and poultry in several Latin American countries [58,59]. Considering these findings and because colistin has been largely used as a growth promotor in Latin America, the poultry industry could be considered an important hotspot for this kind of resistance. Additionally, it has to be considered that although we did not find *mcr-2* in our study, up to 8 genetic determinants for colistin resistance have been described [60-62]. Therefore, a search for more genetic determinants should be conducted to better understand the epidemiology of colistin resistance in poultry from Ecuador.

Genes of the *bla*_CTX-M_ family have been the most prevalent in several studies even when there is no selective pressure due to antibiotic usage [44,63]. In our study, the *bla*_CTX-M_ family was more prevalent than the AmpC genes. A study in Colombia showed that most *E. coli* isolates from poultry had *bla*_CMY_ genes [39]. This study showed that the most common variant of the AmpC gene was *bla*_CMY-2_ (58 isolates with *bla*_CMY-2_ and one *bla*_CMY-46_ isolate).

In our study, *bla*_CTX-M-65_ was the most prevalent allele of the *bla*_CTX-M_ family (33,6% of the ESBL-producing isolates) followed by *bla*_CTX-M-3_ (19,1%) and *bla*_CTX-M-55_ (17,3%) which differs from the results of other countries in the region. Colombia reported *bla*_CTX-M-2_ as the most prevalent variant followed by *bla*_CTX-M-8_ and *bla*_CTX-M-15_ [39], while Brazil identified *bla*_CTX-M-8_ and *bla*_CTX-M-2_ variants in chicken meat [38,40,64] In our case, *bla*_CTX-M-8_ and *bla*_CTX-M-2_ were present as a small proportion. Other genes such as *bla*_SHV-5_, *bla*_SHV-153_ and *bla*_TEM-176_ were found in lower proportions which agrees with the mentioned studies.

In Ecuador, there are no data about *bla*_CTX-M-65_ in *E. coli* from poultry. However, this variant has been identified in *Salmonella* from poultry and in human clinical samples [65]. Likewise, *bla*_CTX-M-3_ and *bla*_CTX-M-55_ have not been reported in Ecuadorian poultry isolates but have been identified in human infections [34,66]. These findings suggest the presence of plasmids carrying these variants in our environment and the possibility of transmission from poultry to human. Finally, 6 isolates did not present ESBL or AmpC genes. In these cases, a broader panel of genes should be used to identify the genetic determinants of resistance in these isolates.

The fact that different ESBL and AmpC genotypes originated from more than 1 farm, suggests that cross contamination between farms is a plausible hypothesis. This idea is supported by other studies that found that *Salmonella* and *Campylobacter* isolated from different poultry farms are clonally related [67]. In Ecuador, climatic and social factors lead to most poultry houses having an open configuration in which implementation of rigorous biosecurity is difficult. Spread of bacterial genotypes among farms and integrated companies can therefore be a common event [68,69]. This highlights the importance of implementing effective biosecurity systems aiming not only to avoid the spread of antimicrobial resistance but also to improve poultry health.

In conclusion, poultry production systems represent a hotspot for antimicrobial resistance in Ecuador, possibly mediated by the extensive use of antibiotics in this industry. Monitoring this sector in national and regional plans of antimicrobial resistance surveillance should therefore be considered.

## Author contributions

Conceived and designed the experiments: CV, LD, JZ.

Performed the experiments: CV, DO, CN.

Analyzed the data: CV, DO, CN.

Contributed reagents/materials/analysis tools: CV, LD, JZ.

Wrote the paper: CV, DO, CN, LD, JZ.

## Acknowledgments

The authors want to thank Pedro Barba and slaughterhouse staff for their technical assistance.

